# Executive control can query hidden human memories

**DOI:** 10.1101/2024.10.22.619676

**Authors:** Chong Zhao, Keisuke Fukuda, Geoffrey F. Woodman

**Affiliations:** Department of Psychology, Center for Integrative and Cognitive Neuroscience, Vanderbilt Vision Research Center, Vanderbilt University, Nashville, Tennessee 37240, USA

## Abstract

When we try to retrieve a representation from visual long-term memory there is a chance that we will fail to recall seeing it even though the memory is stored in our brain. Here we show that although mechanisms of explicit memory retrieval are sometimes unable to retrieve stored memories, that mechanisms of executive control can quickly query memory and determine if a representation is stored therein. Our findings suggest that the representations stored in human memory that cannot be accessed explicitly at that moment are nonetheless directly accessible by the brain’s higher level control mechanisms.

## Introduction

We humans frequently forget that we have previously encountered important things in our environment (Wixted 2022, Wixted & Carpenter 2007), with forgetting becoming more pronounced during aging and clinical disorders (García-Martínez et al 2023). As cognitive psychologists we often use recognition memory tests to access the quality of human memory in the laboratory (Brown & Aggleton 2001). These tasks generally involve presenting to-be-remembered stimuli, such as objects, with later testing requiring subjects to report whether an item is old or new, and their confidence in that behavioral report (**Fig. 1A**). Although visual long-term memory is quite accurate (Brady et al 2008, Shoval et al 2023), it is believed that human memory stores more information than can be explicitly retrieved at any given moment in time (Rugg et al 1998, Voss & Paller 2009). Here we tested the hypothesis that those memories that cannot be explicitly reported in memory experiments can nonetheless be accessed by executive control mechanisms that monitor our behavioral response output and generate distinct electrophysiological signatures.

**Fig. 1.**
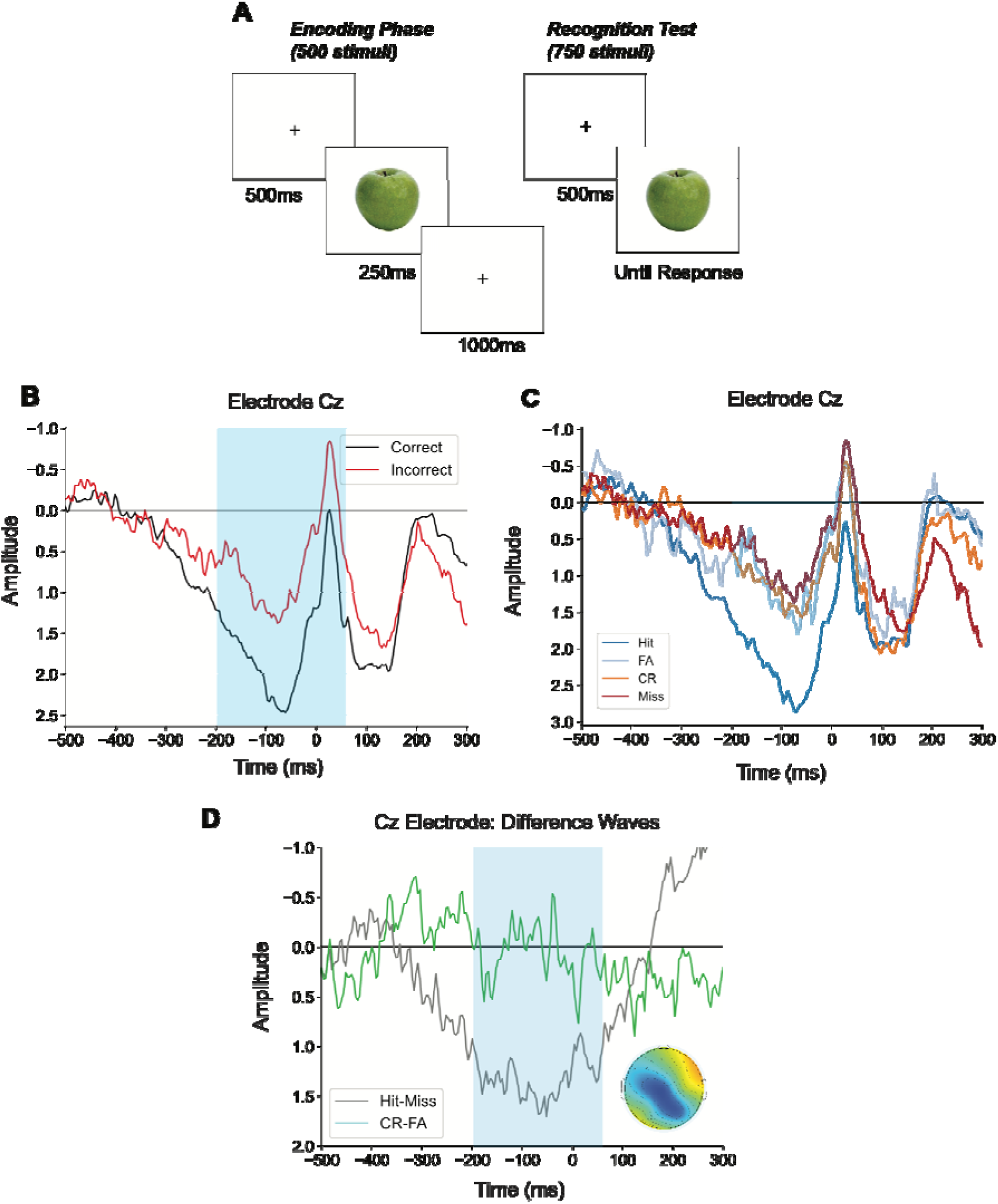
The task our human observers (N=50) performed and the event-related potential results from the electroencephalogram recordings. **A**. An example of the stimuli presented during the encoding and then the test phase of the experiment. B. The waveforms time locked to button press onset for correct and incorrect trials, plotted separately. C. The waveforms sorted by whether the test image was of an old item shown during encoding (hit or miss) or a new item (correct rejection or false alarm). D. The difference waves and scalp distributions of the memory error-related negativity (ERN, measured in the blue time window).

By recording humans’ electroencephalogram (EEG), researchers have used the Error-Related Negativity (or ERN) extensively to study executive control because it is elicited when humans make an error in many different settings (Dehaene et al 1994, Gehring et al 2012, Gehring & Willoughby 2002). For the brain to detect this error, it must compare the behavioral response that was just made to the intended response for that trial. For perceptual-motor tasks it is assumed that the brain codes the intended, correct response (Crapse & Sommer 2008, Nieuwenhuis et al 2001). In the context of a memory experiment, the observation of an ERN has more revealing implications. That is, to elicit this activity associated with error detection following a memory error would show that executive control mechanisms of the brain had access to the necessary memory. The ERN can only be generated if the brain knows whether the test object was previously shown in the memory set. To test this hypothesis we accumulated a large dataset from 50 individuals sampled across two institutions performing the same visual recognition memory task. This large dataset afforded further comparisons of the error-related activity as a function of response speed and confidence.

## Methods

### Participants

We conducted a power analysis to determine the sample size needed for adequate statistical power, employing an effect size of 0.5 and a significance level of 0.05. The analysis revealed that a minimum of 34 subjects would be necessary to achieve a power of 0.8, while a sample size of 44 subjects would be required to attain a power of 0.9. Subjects were recruited from Vanderbilt University and the University of Toronto and paid for their participation. For the Vanderbilt sample, 23 participants (10 males and 13 females, aged 18–32) took part after providing informed consent, with all procedures approved by Vanderbilt University’s Institutional Review Board. Participants were compensated with $30. All participants self-reported being neurologically healthy, with normal or corrected-to-normal vision, and no color vision deficiencies. Data from three individuals were excluded due to incomplete session participation. After artifact rejection, 19 subjects were kept in our final analyses.

For the University of Toronto sample, thirty-eight young adults (22 females; M = 20.32 years old, S.D. = 2.55, Range = 18-32) from the University of Toronto Mississauga community participated in return for monetary compensation (12 CAD/hour). Every participant provided written informed consent to the protocol approved by the Research Ethics Boards of the University of Toronto. Data obtained from 7 participants were rejected due to excessive EEG artifacts (> 40% trials rejected due to artifacts), resulting in 31 participants in the initial analysis. In total, our experiment contained 50 quality subjects for our final analysis.

### Stimuli and Procedures

The stimuli and task setup are depicted in **Fig. 1A**. The stimuli were adapted from an established set of photographs (Brady et al 2008). During the encoding task, participants were shown 500 sequential images of real-world objects, with brief pauses every 50 images. Participants were instructed to study each image while maintaining central fixation, preparing for a subsequent recognition-memory test. Each trial began with a button press on a gamepad, followed by a 1250 ms pre-encoding interval, a 250 ms presentation of the image, and then a 1000 ms blank screen during encoding. A central fixation dot then signaled the start of the next trial. After the encoding phase, participants’ resting-state EEG activity was recorded for 15 minutes with eyes open and closed. Memory for the images was assessed using a recognition test.

The recognition memory test commenced with the appearance of a central fixation dot. Participants initiated each trial with a button press and were instructed to maintain central fixation without blinking throughout the trial. Following a 1250 ms blank screen, an image of a real-world object was displayed at the center of the screen, with a mix of new and previously seen images presented in random order. After the 1250 ms image presentation, a blue and a red dot appeared, each on opposite sides of the image. Participants then indicated whether they remembered seeing the image during the study phase. The red dot’s position indicated which side of the gamepad to press if the image was remembered, while the blue dot indicated the buttons to press if it was not. Participants used a scale to indicate their confidence, pressing different buttons for 100%, 80%, or 60% confidence. The red and blue dot positions were randomized in each trial. After responding, participants had a self-paced break to rest and blink. They were tested on 500 previously seen images and 250 new images.

### EEG Acquisition and Pre-processing

EEG recordings were made using a right-mastoid reference, with offline re-referencing to the average of both mastoids. Electrodes were placed at standard 10–20 system locations (Fz, Cz, Pz, F3, F4, C3, C4, P3, P4, PO3, PO4, O1, O2, T3, T4, T5, and T6) and at custom locations OL (halfway between O1 and OL) and OR (halfway between O2 and OR). Eye movements were tracked with electrodes placed 1cm lateral to the external canthi for horizontal movements and beneath the right eye for vertical movements and blinks. Signals were amplified with a gain of 20,000, bandpass filtered between 0.01–100 Hz, and digitized at 250 Hz. Trials with horizontal eye movements exceeding a 30µV threshold or eye blinks exceeding a 75µV threshold were excluded from further analysis.

### ERP Analyses

To examine the ERPs associated with memory test response, data were time-locked to the button press that indicated response during each trial, analyzing waveforms from -500ms to +300ms relative to the motor response. To observe the ERN emerge around the time of response we baseline-corrected each EEG segment to the mean amplitude from -500ms to -300ms before the measurement epoch of interest. We entered the mean amplitudes measured from -200 to 50 ms around the time of the button press into an ANOVA with the factors of correctness (correct versus error) and electrode (Fz, Cz, versus Pz), with subsequent analyses examining median splits based on each individual’s median RT or confidence rating across all stimuli. We also analyzed our data for the error positivity (or Pe) which follows the ERN and is maximal over parietal electrodes. We did not find that the positivity following the ERN was significantly modulated by the status of the test object, unlike the ERN. In **Figure S1**, we show the FN400 waveforms. The FN400 or frontal old/new effect appears to measure the familiarity, implicit memory, or perceptual fluency with which a stimulus is processed (Voss & Federmeier 2011). We show these waveforms to verify that the pattern of effects across the ERN are different than those of the FN400, which overlaps it in time.

## Results

As shown in **Fig. 1B and 1C**, we found that subjects’ waveforms were more negative just as they were about to commit an error. The ERN is defined as a more negative potential around the time of a behavioral response on error trials relative to trials with correct responses (Fz: *t*(49) = 2.38, *p* = 0.02, Cohen’s *d* = 0.34; Cz: *t*(49) = 3.37, *p* = 0.001, Cohen’s *d* = 0.48). This effect began approximately 200 ms before the button press was made and continued for approximately 100 ms afterward. It is known that the ERN can precede the commission of the erroneous behavioral response, presumably because the difference between the actual processing state and the intended processing state is detected prior to the final stages of response execution (Falkenstein et al 2000, Rodriguez-Fornells et al 2002). The scalp distribution of this effect is similar to previous studies of the ERN using other tasks (**Fig. 1E blue inset**)(Dehaene et al 1994, Reinhart et al 2012). Inferential statistical tests showed that error trials elicited more negative potentials relative to correct trials evidenced by a main effect of accuracy (*F*(1,49) = 11.56, *p* =0.001, η_p_^2^ = 0.13) in the ANOVA of voltage, and because this difference was larger in amplitude at central relative to frontal electrodes we observed an interaction of electrode x trial type (*F*(1,49) = 7.68, *p* =0.008, η_p_^2^ = 0.1, due to large ERN at Cz than Fz).

Our next goal was to determine the relationship between the ERN and the nature of the behavioral responses that it is time-locked to. Our first relevant observation is that the ERN exhibited a large amplitude difference between miss trials and hit trials (*t*(49) = 2.98, *p* = 0.004, Cohen’s *d* = 0.42), but there was not a significant difference in the response-locked waveforms between false alarm and correct rejection trials (*t*(49) = 0.08, *p* = 0.93, see **Fig. 1C**). This observation of an ERN elicited on miss trials relative to hit trials (i.e., when old test items are shown), but not false alarms relative to correct rejections (i.e., when new test items are shown) indicates that the executive control mechanism that generates the ERN can detect when an test item matches a representation in memory, but does not return diagnostic evidence about the behavioral response being inaccurate when no memory match can be found. This result shows an parallel to other cognitive tasks in which observers need to determine whether a given object is absent from a visual or memory set (Chun & Wolfe 1996, Treisman & Gormican 1988, Zhao et al 2023). Next, we analyzed our data to determine whether this error-related activity was sensitive to the speed or confidence of the observers’ judgements about whether they remembered something.

To determine with the memory-related ERN was related to the speed of subjects’ behavior, we analyzed the test stimulus locked ERPs based on accuracy and divided by median response time, the ERN amplitude was found to be more negative for incorrect trials compared to correct trials (Main Effect of accuracy: *F*(1,49) = 4.75, *p* = 0.03, η_p_^2^ = 0.10) and more negative for slower trials compared to faster ones (Main Effect of RT: *F*(1,49) = 5.42, *p* = 0.02, η_p_^2^ = 0.11, see **Fig. 2A**). In line with the results in **Fig. 1**, fast hit and fast miss trials exhibited a significant different in amplitude between each other (*t*(49) = 2.57, *p* = 0.01, Cohen’s *d* = 0.38), whereas fast correct rejection and fast false alarm trials did not (*t*(49) = 0.29, *p* = 0.77, see **Fig. 2B**). This suggests that a more negative ERN amplitude was associated with a more challenging retrieval process during the test phase, resulting in longer response times.

**Fig. 2.** Results showing relationships between subjects’ behavior and the error-related negativity (or ERN). **A**. The ERN plotted as a function of accuracy and RT at electrode Cz. **B**. The ERN as a function of response speed, using a median split. **C**. The ERN as a function of response speed and the type of test item presented.

Presumably, executive control mechanisms can query both explicitly available and currently unavailable representations from memory, such that the amplitude of this metric should be correlated with subjective reports, as with the RT analyses that we just discussed. **Fig. 2C** shows that the waveforms that exhibited a significant effect of correctness (*F*(1,49) = 7.44, *p* = 0.009, η_p_^2^ = 0.13), and a significant interaction of electrode and confidence (*F*(1,49) = 5.69, *p* = 0.02, η_p_^2^ = 0.10) due to the modulation by confidence being maximal at location Cz, the electrode shown in **Fig. 2A-C**. These findings suggest that the pattern of ERN activity was sensitive to the confidence that subjects had when they made their behavioral responses, as we would expect if this mechanism of executive control can access all the contents of memory.

In classic studies of the ERN, researchers reported that its amplitude is related to the degree to the size of the compensatory behavior effects found on the following trial (Gehring et al 1993). In the present study, subjects reported when they were ready for another memory test by initiating the next test with a space bar press. Our findings appear unconclusive as yet, because we lacked sufficient resolution to observe a significant slowing of initiation time following a hit compared to a miss (*t*(49) = 1.72, *p* = 0.09), although such a trend was observed, with no such slowing between correct rejections and false alarms (*t*(49) = 1.07, *p* = 0.29), mirroring the pattern of medial-frontal negativities on these trials. We believe that it would be fruitful for subsequent studies to focus on determining whether error detection during testing memory results in compensatory behavior on the next event.

## Discussion

We observed a medial-frontal negativity around the time of our subjects’ button presses that was more negative for error responses than for correct responses during visual long-term memory recognition testing. This medial-frontal negativity had a scalp distribution that was similar to that observed in previous studies of the ERN and was related to the speed and confidence of those trial-by-trial recognition judgements.

The present empirical observations have important theoretical and practical implications. Theoretically, the present findings show that high-level executive control mechanisms appear to be able to access the memory representations that are stored in the human brain, apparently via a route that is independent of the explicit retrieval mechanisms. Although it has been known that the human brain can implicitly access memory representations that cannot be explicitly retrieved by performing tasks with greater efficiency across experience (Reber 2013, Roediger 1990, Rugg et al 1998), the present paper shows for the first time that the brain has mechanisms that can quickly query those memories, even those memories cannot be explicitly accessed to guide behavior at that moment. This counterintuitively suggests that executive control mechanisms may have memory access that even memory mechanisms lack.

Practically, the present finding suggest that it may be possible to develop brain-computer interfaces that can detect when the user has a memory stored, even when they fail to report it. That is, in principle an algorithm trained on a person’s own medial-frontal negativities could probe the depths of memory storage to determine whether that individual has a currently hidden memory. Clearly the ethical use of this kind of memory readout ability will require more extensive discussion in broader society.

## Acknowledgements

We thank Gordon Logan, Sisi Wang, and Sean Polyn for formative discussions of this work. This work was made possible by grants from the National Science Foundation (BCS-2147064), and the National Institutes of Health (P30-EY08126 and T32-EY007135).

---

**Fig. S1.**
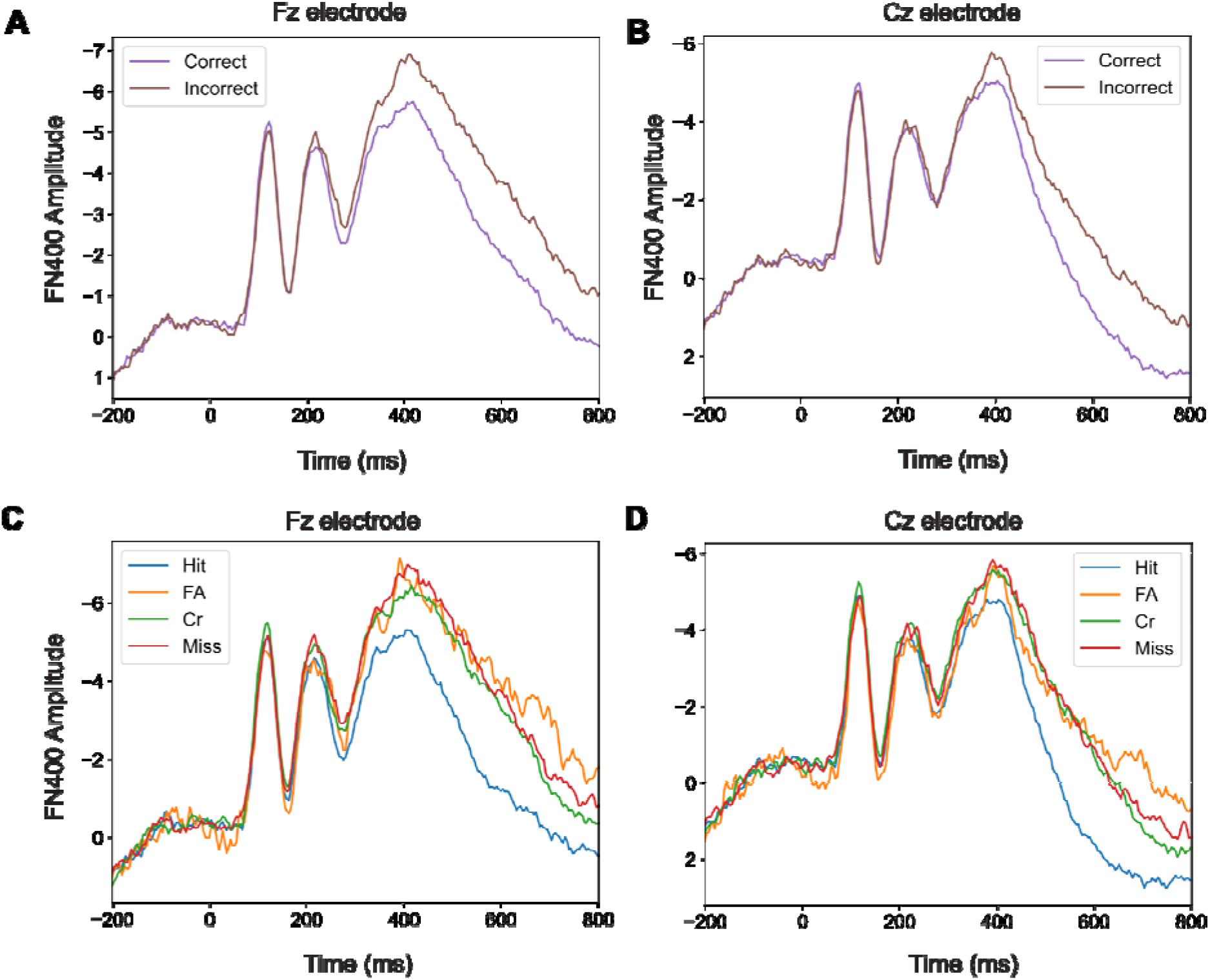
The analyses of additional ERP waveforms time locked to the test stimulus onset. **A**. The FN400 or frontal old/new effect as a function of whether the subject would get the trial correct or not. B shows that the scalp distribution of the FN400 was frontal, maximal at Fz. C. The waveforms sorted by whether the test image was of an old item shown during encoding (hit and miss) or a new item (correct rejection or false alarm). D. The same waveforms shown at electrode CZ.

